# Comparison of the cattle rumen bacterial community and drinker water in different livestock farm buildings using NGS sequencing

**DOI:** 10.1101/2023.08.21.554094

**Authors:** E.A. Yildirim, L.A. Ilina, V.A. Filippova

## Abstract

The aim of the study was to analyze samples of ruminal fluid from highly productive lactating cows of black-motley Holsteinized breed and water from drinkers from three livestock farm buildings using the molecular genetic method NGS -sequencing. The experiment was carried out in a commercial farm of the Tula region of Russia. The composition of microorganisms in samples of rumen fluid and water from drinking bowls was studied using the molecular genetic method of NGS-sequencing. In all samples, including water samples from drinking bowls, a variety of pathogenic microorganisms were found, including campylobacter, mycoplasma, pathogenic Clotridia, leptotrichia, Salmonella and others. Interestingly, a number of these microorganisms were also found in the rumen of cows. Thus, the proportion of pathogenic Campylobacter in samples of cicatricial contents ranged from 0.01±0.0006% to 0.03±0.0009%. Also, these bacteria were found in water samples from the drinking bowls of two buildings: in the amount of 0.02±0.0015% and 0.002±0.0001%.Improper sanitary condition of water in cowsheds creates the risk of various diseases.

## Introduction

Scientifically unjustified feeding leads to undesirable changes in the rumen microbiocenosis, which is the cause of a decrease in productivity and the occurrence of a number of diseases due to a violation of the digestive processes. Under conditions of intensive animal husbandry, the composition of the environment, feed and water has a direct impact on the qualitative and quantitative characteristics of the microbial community of the gastrointestinal tract of animals [1]. Improper sanitary condition of cowsheds creates the risk of various diseases that can disrupt the rhythm of production and lead to economic losses. A significant factor influencing the development of agricultural enterprises is the quality of water, which is an integral part of the habitat of most living organisms.

Some information about the composition and role of microflora was obtained using classical methods of microbiology. However, these methods have a number of significant limitations and disadvantages, including the impossibility of correct counting of microbes due to their growth on nutrient media for cultivation not from one cell, but from their accumulation. The emergence and development of modern molecular genetic methods has made it possible to study the diversity of microorganisms without the limitations associated with traditional methods of microbiology - i.e. bypassing the stage of cultivation [2].

One of the most promising today is NGS -sequencing (English next generation sequencing). The use of next-generation sequencing technologies makes it possible to carry out metagenomic studies of complex microbial communities with a larger volume of read nucleotide sequences than when using Sanger sequencing. In this case, it is possible to accurately determine the phylogenetic affiliation of microorganisms to a species.

### The aim of the study

was to analyze samples of ruminal fluid from highly productive lactating cows of black-motley Holsteinized breed and water from drinkers from three livestock farm buildings using the molecular genetic method NGS -sequencing.

## Materials and methods

The experiment was carried out in a commercial farm under the conditional number “1” of the Tula region of Russia. Cows of black-motley Holsteinized breed of the 2nd and 3rd lactation with a productivity of 10,000 kg were kept in three different livestock farm buildings. The live weight of cows was 600 kg. The animals were kept under the same conditions. The content of animals is tethered. Cow rations were calculated automatically using the program “AMTS.Cattle.Professional” (https://agmodelsystems.com) in accordance with generally accepted requirements. Groups were formed - analogues: 3/1, 4/1, 4/4, 3 animals in each group, which were kept in different buildings. From drinkers of groups 3/1, 4/1, 4/4 were taken water samples in triplicate.

Sampling of the contents of the rumen of cows (in triplicate) was carried out from the upper part of the ventral sac of the rumen of cows with the maximum possible compliance with aseptic conditions with this method manually using a sterile probe. The selected samples of the rumen and water were immediately placed in sterile plastic tubes. All samples were frozen at – 20°C and transported in dry ice to the laboratory for subsequent DNA isolation.

Total DNA was isolated from the samples using the Genomic DNA Purification Kit (Thermo Fisher Scientific, Inc., USA) according to the attached instructions. The bacterial community was assessed by NGS-sequencing, on an automatic sequencer MiSeq (Illumina, Inc., USA) using primers for the V3–V4 region of 16S rRNA: 5’ — TCGTCGGCAGCGTCAGATGTGTGTATAAGAGACAGCCTACGGGNGGCWGCAG — 3’ (forward primer), primer). PCR conditions were as follows: 3 min at 95°C; 30 s at 95°C, 30 s at 55°C, 30 s at 72°C (required for sequence extension) (25 cycles); 5 min at 72°C (final extension). Sequencing was performed using Nextera ® XT IndexKit library preparation reagents (Illumina, Inc., USA) and Agencourt PCR products for purification. AMPure XP (Beckman Coulter, Inc., USA) and for sequencing MiSeq® ReagentKit v2 (500 cycle) (Illumina, Inc., USA). The maximum length of the resulting sequences was 2 × 250 bp.

Automatic bioinformatic data analysis was performed using QIIME2 ver software 2020.8 (https://docs.qiime2.org/2020.8/). After importing sequences in fastq from the sequencing instrument and creating the necessary mapping files (containing the metadata of the files being studied), paired read lines were aligned. Next, the sequences were filtered for quality using the default settings. Noise sequences were filtered using the DADA2 package built into the QIIME2 software, which includes information about the quality of sequences in its error model (filtering of chimeric sequences, artifacts, adapters), which makes the algorithm resistant to a sequence of lower quality. In this case, the maximum length of the pruning sequence was used, equal to 250 bp (https://benjjneb.github.io/dada2/tutorial.html). To build a phylogeny de novo performed multiple sequence alignment using the MAFFT software package, followed by masked alignment to remove positions that differed significantly. The taxonomy was assigned using the QIIME2 software, which assigns sequences a taxonomic identification based on ASV data (using BLAST, RDP, RTAX, mothur and uclust methods) using a 16s rRNA database. Silva 138.1 (https://www.arb-silva.de/documentation/release-138.1/).

Mathematical and statistical processing of the results was carried out by the method of multivariate analysis of variance (multifactor ANalysis Of VAriance, ANOVA) in Microsoft Excel XP/2003, R-Studio (Version 1.1.453) (https://rstudio.com). Significance of differences was established by Student’s t-test, differences were considered statistically significant at P≤0.05. Means were compared using the Tukey Significantly Significant Difference (HSD) test and the TukeyHSD function in the R Stats package.

## Research results

The study showed that in the samples of ruminal fluids of cows from buildings No. 3/1 and No. 4/1, the main proportion of microorganisms were representatives of the phyla *Firmicutes* and *Bacteroidetes*, which include, among other things, many representatives of the normoflora that contribute to the processes of digestion of feed [3]. Also, in the studied samples, the presence of the phylum *Proteobacteria* was noted, which includes many pathogenic species.

In samples from cows from housing No. 4/4, a high proportion (up to 82 ± 4.5%) of the entire rumen microbiota was occupied by uncultivated bacteria.

In water samples, non-culturable bacteria also occupied the main share.

As can be seen from figure 2, among the main representatives of cellulolytic bacteria, the order *Bacteroidales*, families *Eubacteriaceae, Flavobacteriaceae, Lachnospiraceae, Ruminococcaceae* and *Clostridiaceae* were found. Cellulolytic bacteria are the most important bacteria that contribute to the breakdown of fiber in plant feeds. The content of this group of microorganisms in the rumen indicates the ability of the animal to assimilate feed rations.

**Fig. 1.**
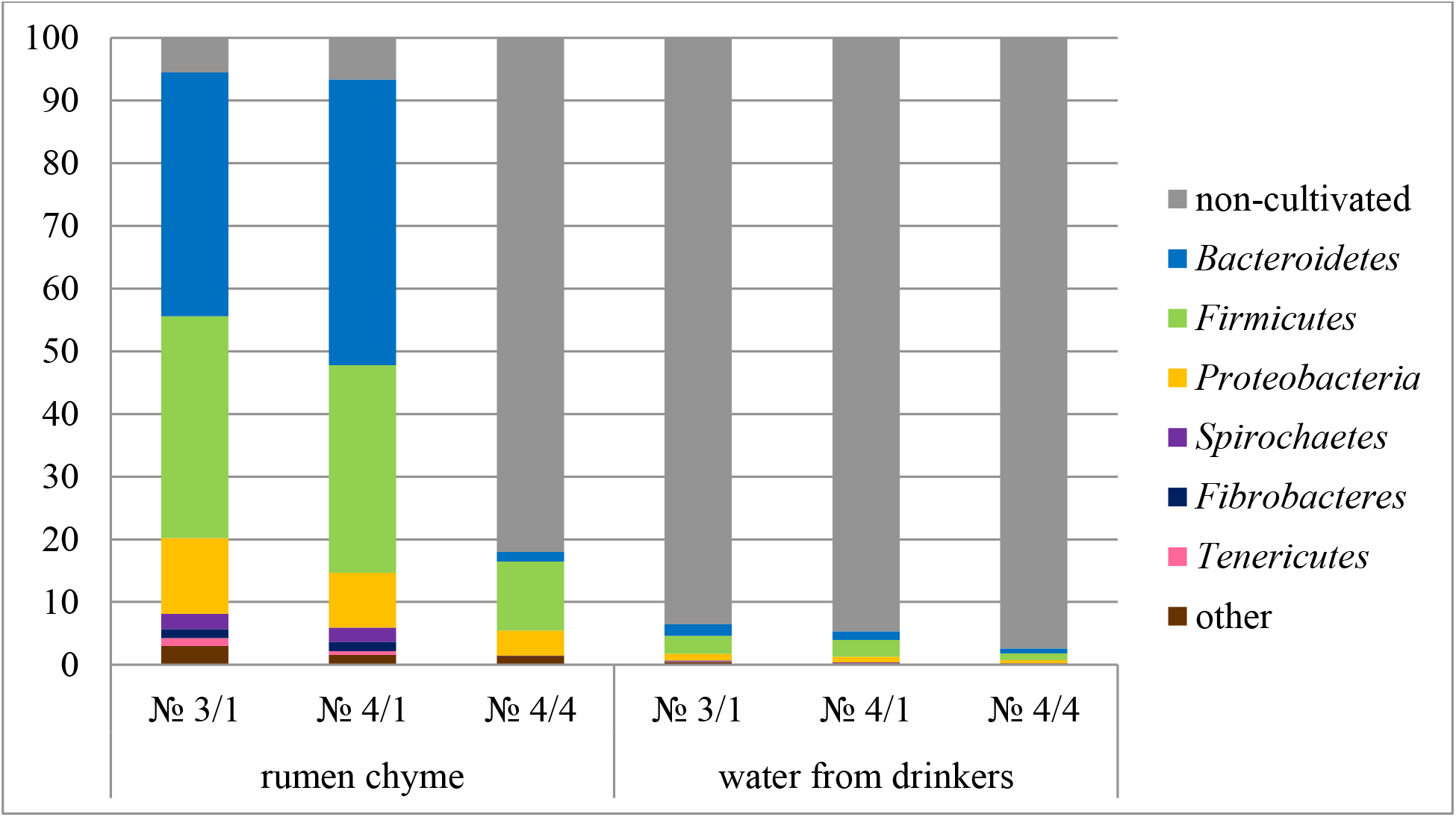
Microbial landscape of the studied samples at the phylum level (according to NGS - sequencing analysis), %

**Fig. 2.**
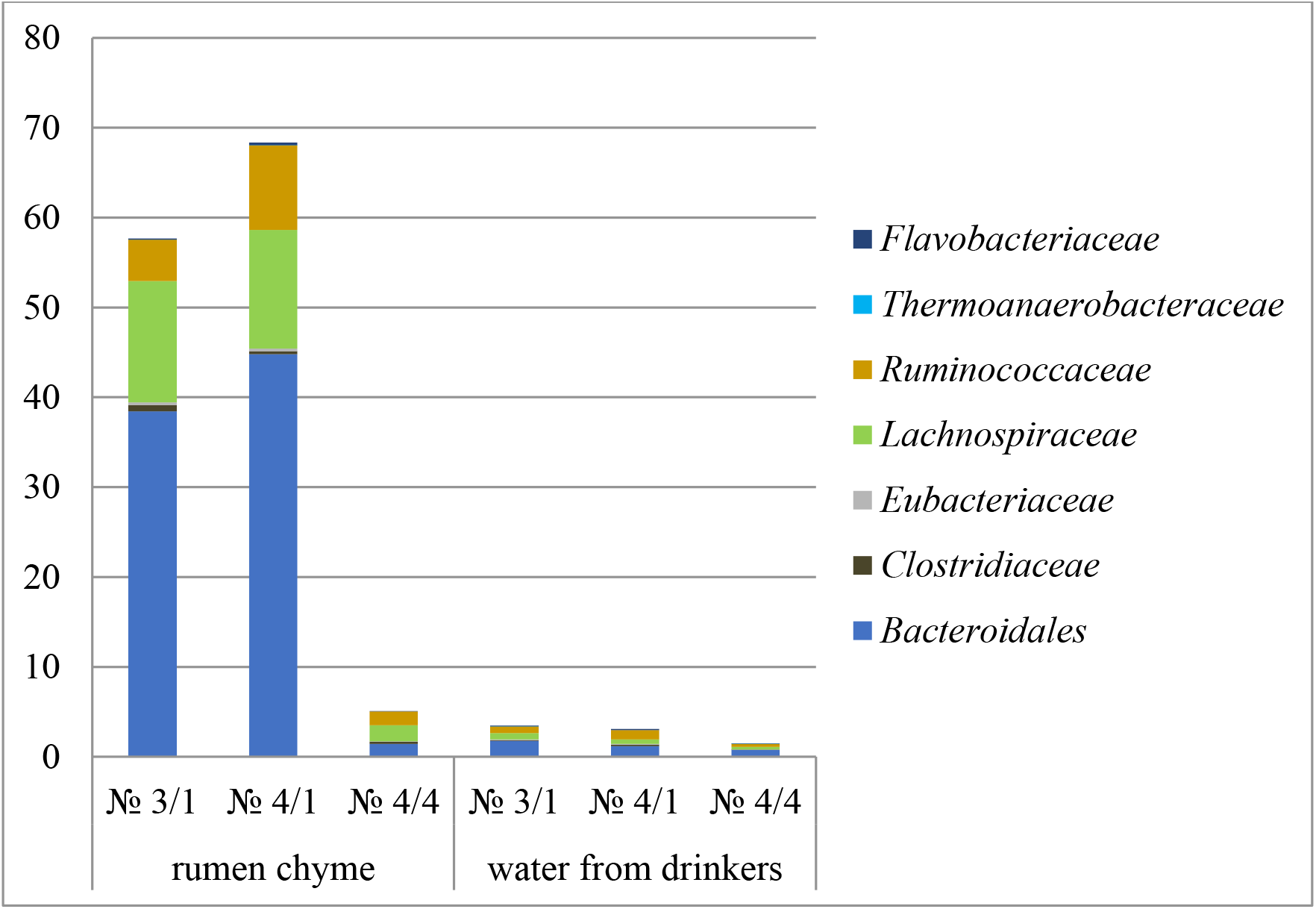
The content of cellulolytics in samples of the ruminal fluid of cows and water from drinkers (according to the analysis by NGS-sequencing), %.

**Fig. 3.**
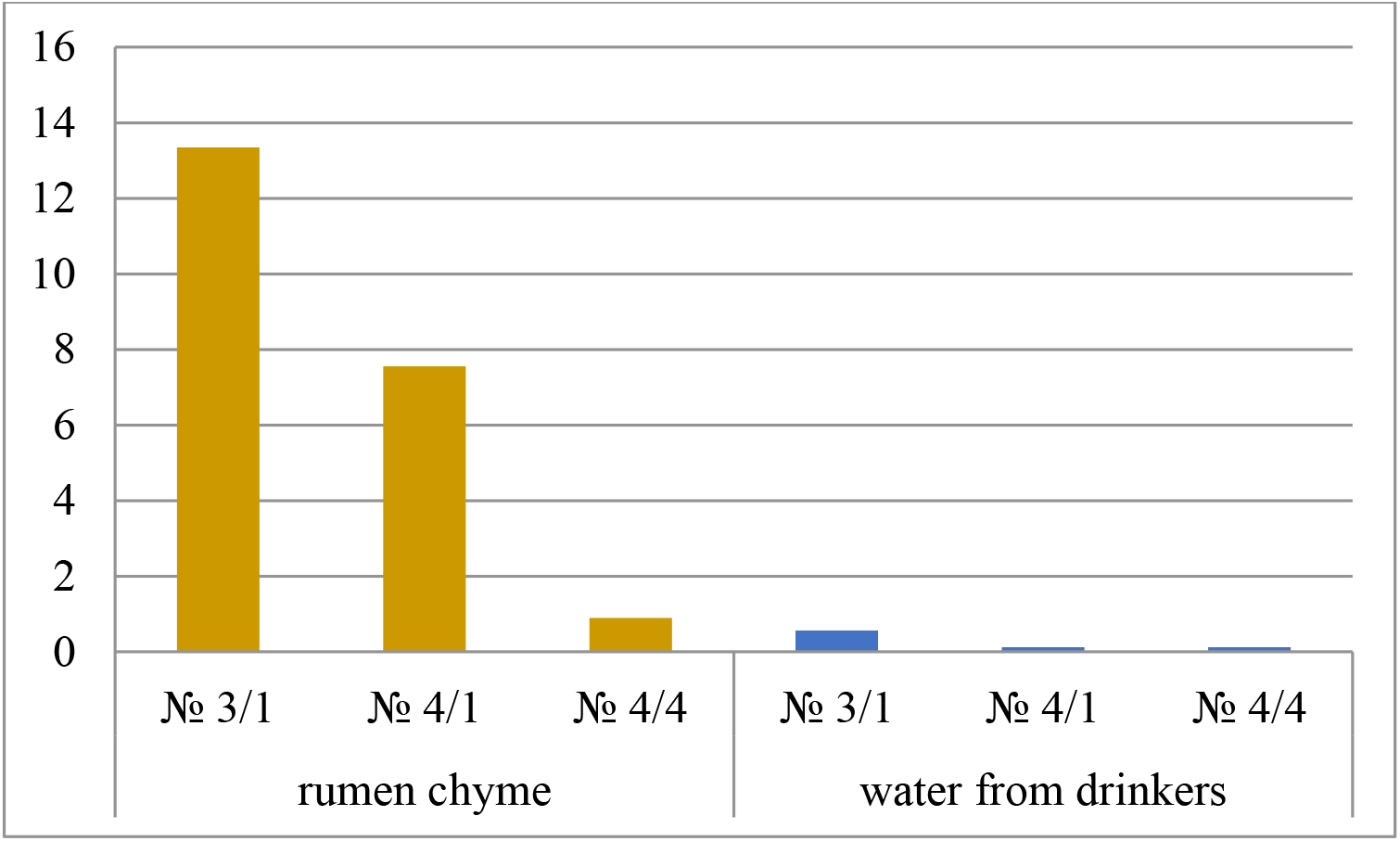
Content of VFA-synthesizing bacteria (according to analysis by NGS sequencing), %

The proportion of cellulolytic microorganisms was high in samples of the contents of the rumen of cows from buildings No. 3/1 (57.7±3.94% in total) and No. 4/1 (up to 68.3±4.56% in total). However, in samples from building No. 4/4, the proportion of cellulolytic microorganisms was not more than 5±0.33% (28.3% of the number of identified species).

Analysis of the samples showed a high proportion of selenomonads in the sample of the content of the cattle rumen from the farm building No. 3/1 - 13.4±0.72%. The proportion of selenomonads was also relatively high in samples of the contents of the rumen of cows from building No. 4/1 - 8.1 ± 0.50%. In samples of cicatricial contents from farm building No. 4/4, the proportion of selenomonads was low, no more than 0.9 ± 0.05% (4.9% of the number of identified species). VFA-synthesizing bacteria (selenomonads) ferment lactic acid produced by bacterioids and lactic acid bacteria (as well as other organic acids) to volatile fatty acids used by the cow’s organism in metabolic processes.

The total content of bifidobacteria (Fig. 4) was relatively low. It is interesting that the highest proportion of bifidobacteria was found in samples of cicatricial contents from farm building No. 4/4 - 0.25±0.017% (1.4% of the number of identified species). In other samples, including water samples, the percentage of bifidobacteria did not exceed 0.13±0.008%. Functions of bifidobacteria in the rumen of cows: antimicrobial activity, immunomodulatory activity, synthesis of vitamins, synthesis of some essential amino acids.

**Fig. 4.**
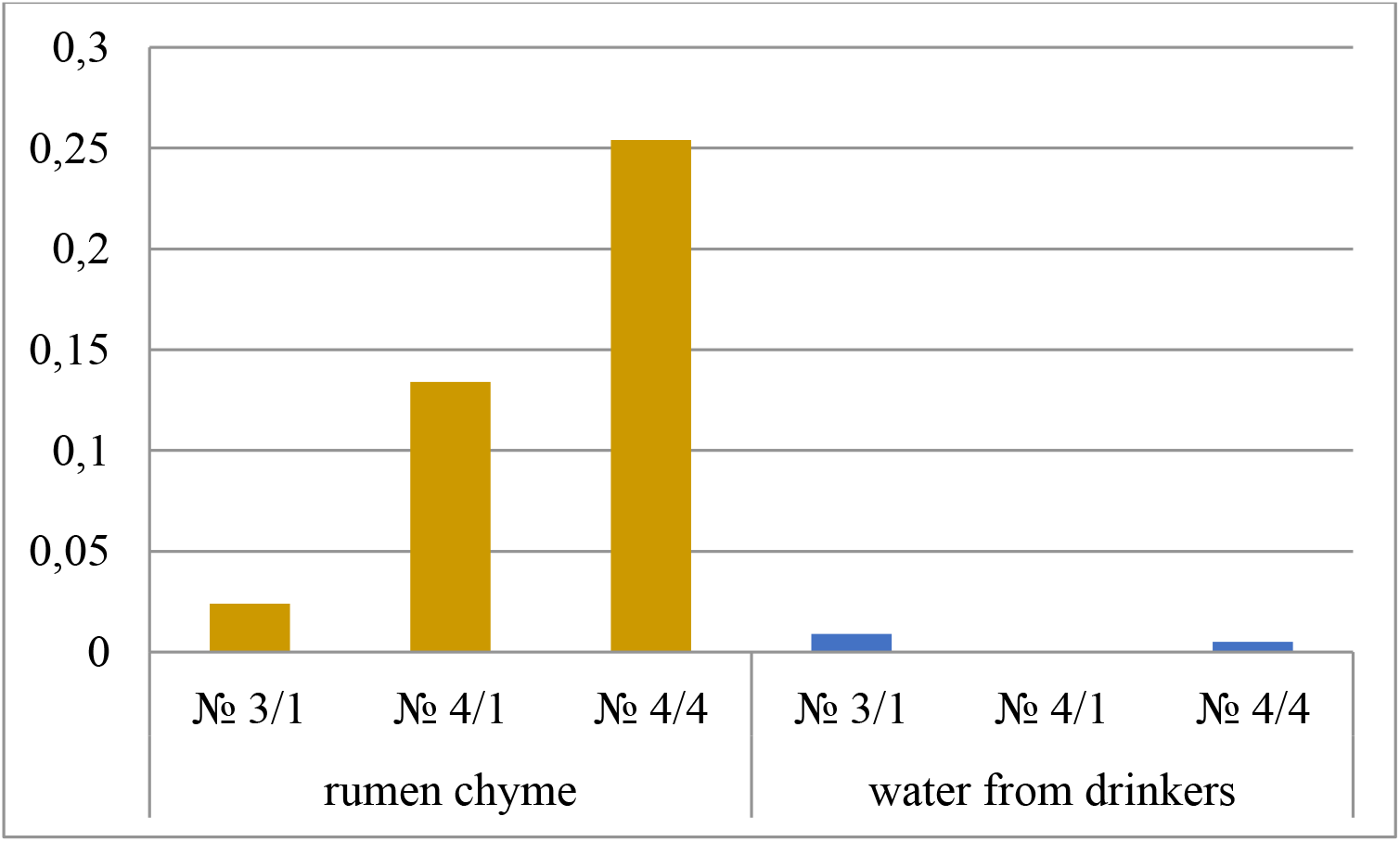
The content of bifidobacteria (according to the analysis by NGS-sequencing) (according to NGS-sequencing analysis), %

Opportunistic microorganisms are often found in the digestive system of healthy animals, but when the organism is weakened, they can multiply rapidly and cause various diseases.

The highest content of opportunistic microbial species was found in samples of cicatricial contents from building No. 4/4 - the total content was 2.3 ± 0.13% (12.9% of the number of identified species), and the main part (2.0 ±0.15%, 11.2% of the number of identified species) were representatives of the genus *Escherichia/Shigella*, including *Escherichia/Shigella dysenteriae*, the causative agent of dysentery (Fig. 5). Also in this sample, 0.16 ± 0.012% (0.86% of the number of identified species) of bacteria of the genus *Corynebacterium* were found, which were not found in other samples of cicatricial contents. Among the identified representatives of this genus, pathogens of bacteremia, inflammation of the oral cavity, pathogens of ulcers were found.

**Fig. 5.**
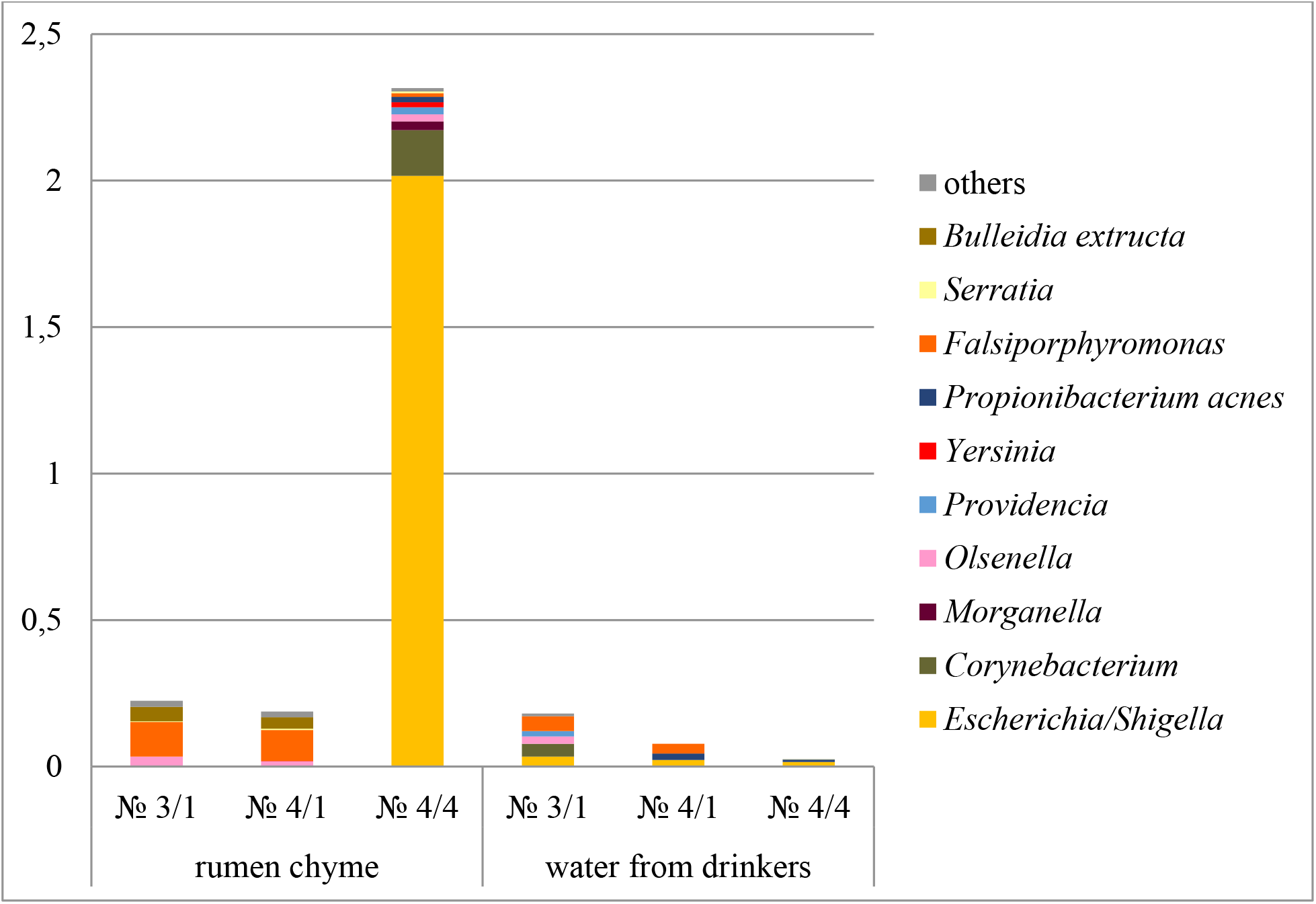
The content of representatives of opportunistic bacteria (according to the analysis by NGS-sequencing), %

The content of representatives of opportunistic microflora in other samples was significantly lower and did not exceed 0.22±0.024% in ruminal fluid samples and 0.08±0.004% in water samples.

As can be seen from Figures 6-8, pathogenic species were detected in all samples studied. The highest content of pathogenic microbial species was found in samples of cicatricial contents from building No. 4/4 - the total content was 1.9 ± 0.12% (10.8% of the number of identified species), and more than half (1.0 ± 0 06%, 5.8% of the *number* of identified species) were pathogenic representatives of the genus *Enterococcus*: *E. faecalis, E. casseliflavus, E. cecorum, E. gallinarum*. Although the latter two species of enterococci are more common as pathogens in birds, their detection in the ruminal fluid of cows is also alarming.

Representatives of streptococci and staphylococci, the causative agents of septic-necrotic diseases, were found in all the studied samples of cicatricial fluids and water from drinking bowls (Fig. 6). Their largest proportion was found in samples of rumen fluid from cows from building No. 4/4: 0.3±0.02% of staphylococci (1.64% of the number of identified species) and 0.3±0.03% of streptococci (1, 64% of the number of identified species).

**Fig. 6.**
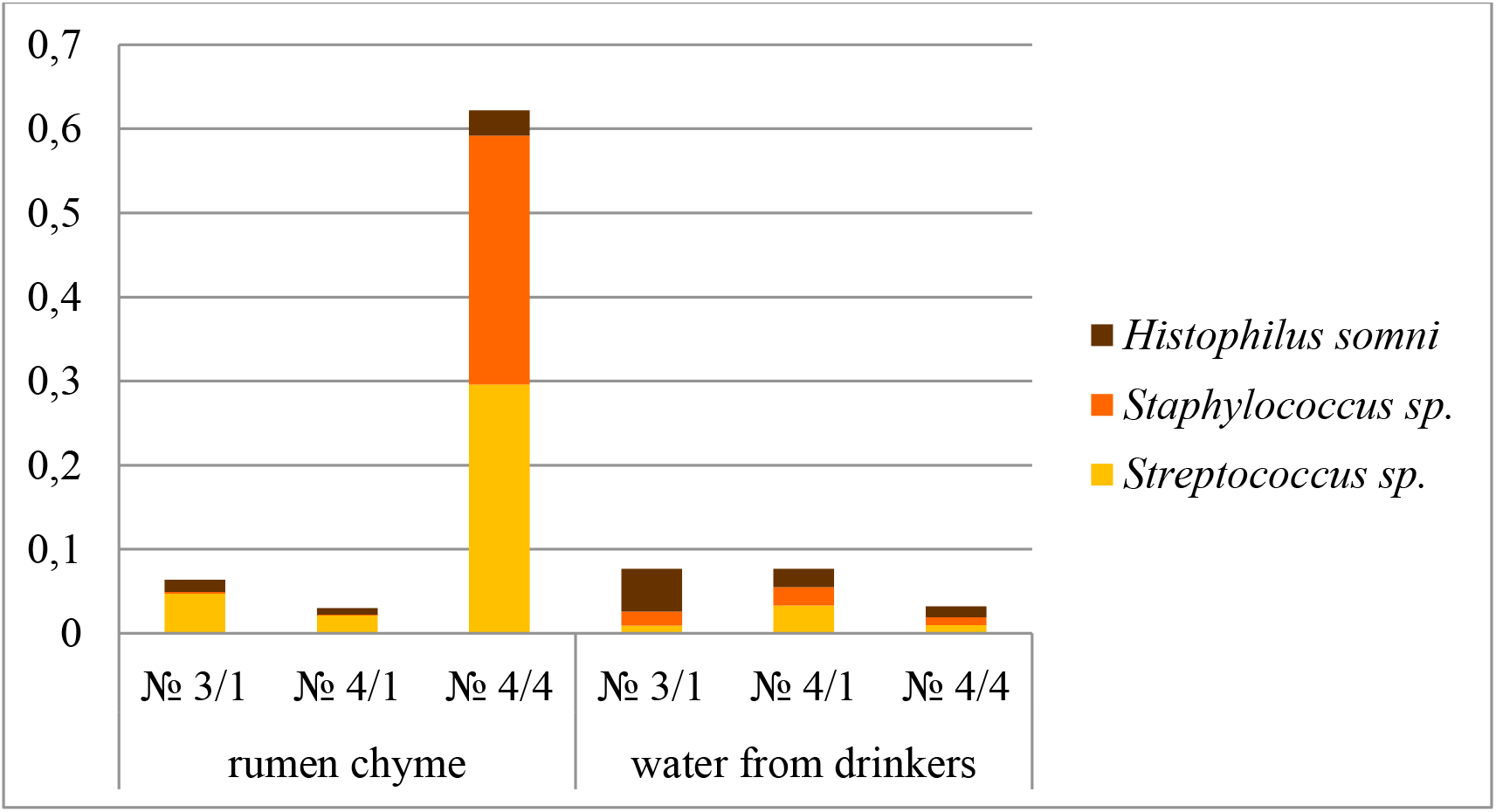
The content of pathogenic streptococci, staphylococci in samples, as well as the species *Histophilus somni* (according to NGS-sequencing analysis), %.

Also, in all the studied samples of rumen fluids and water from drinking bowls, a pathogenic species *Histophilus somni* was found. It often causes inflammation of various organs and systems in livestock (Fig. 6). It is noteworthy that in water samples the proportion of this pathogen was higher (0.01±0.0005% - 0.05±0.003%) than in ruminal fluid samples (0.008±0.0005% - 0.03±0.002%).

In all samples of the contents of the ruminal fluid and in some of the water samples, pathogenic representatives of the genera *Porphyromonas, Campylobacter* and *Mycoplasma* were found (Fig. 7).

**Fig. 7.**
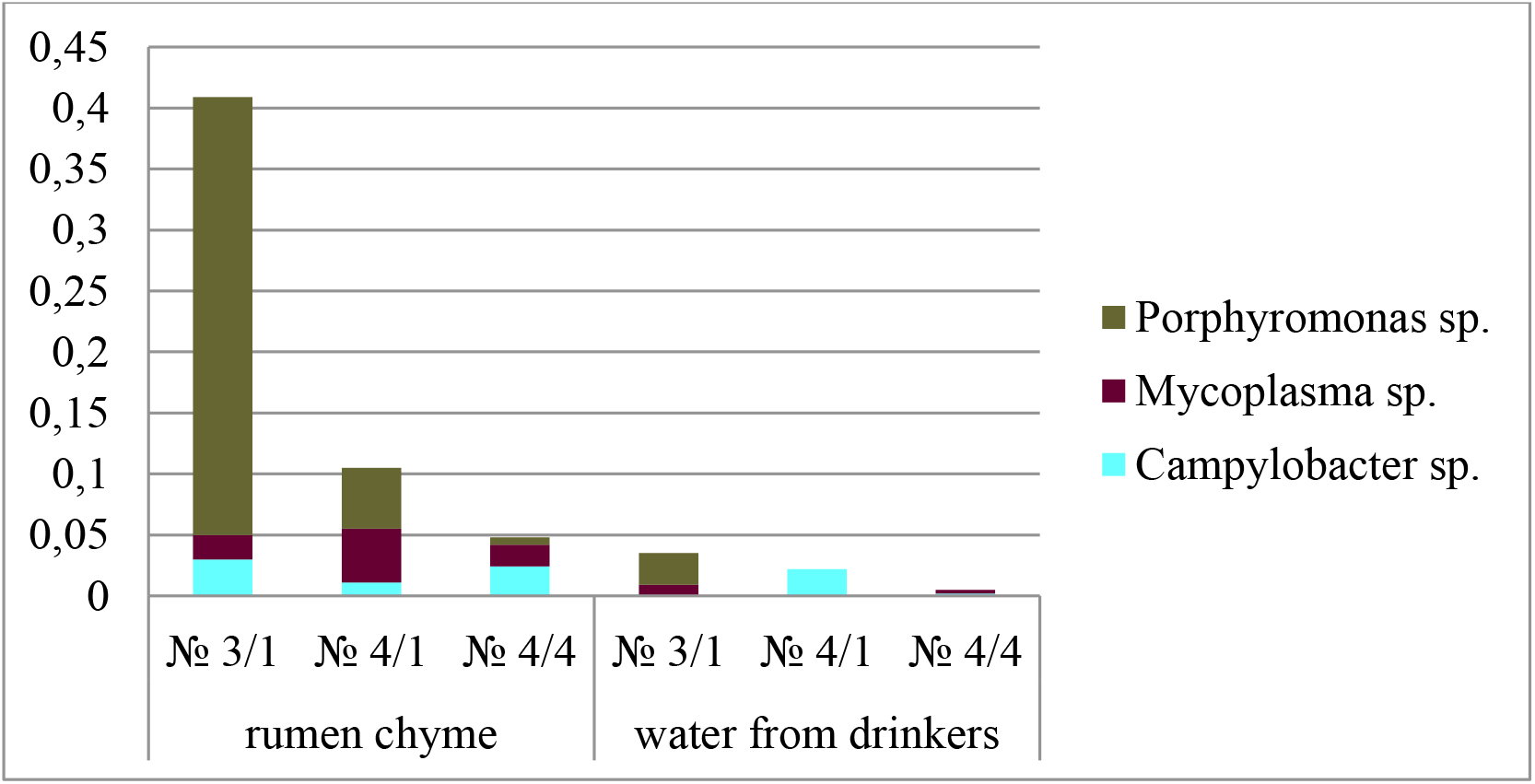
The content of pathogenic representatives of the genera *Porphyromonas, Campylobacter* and *Mycoplasma* (according to NGS-sequencing analysis), %

**Fig. 8.**
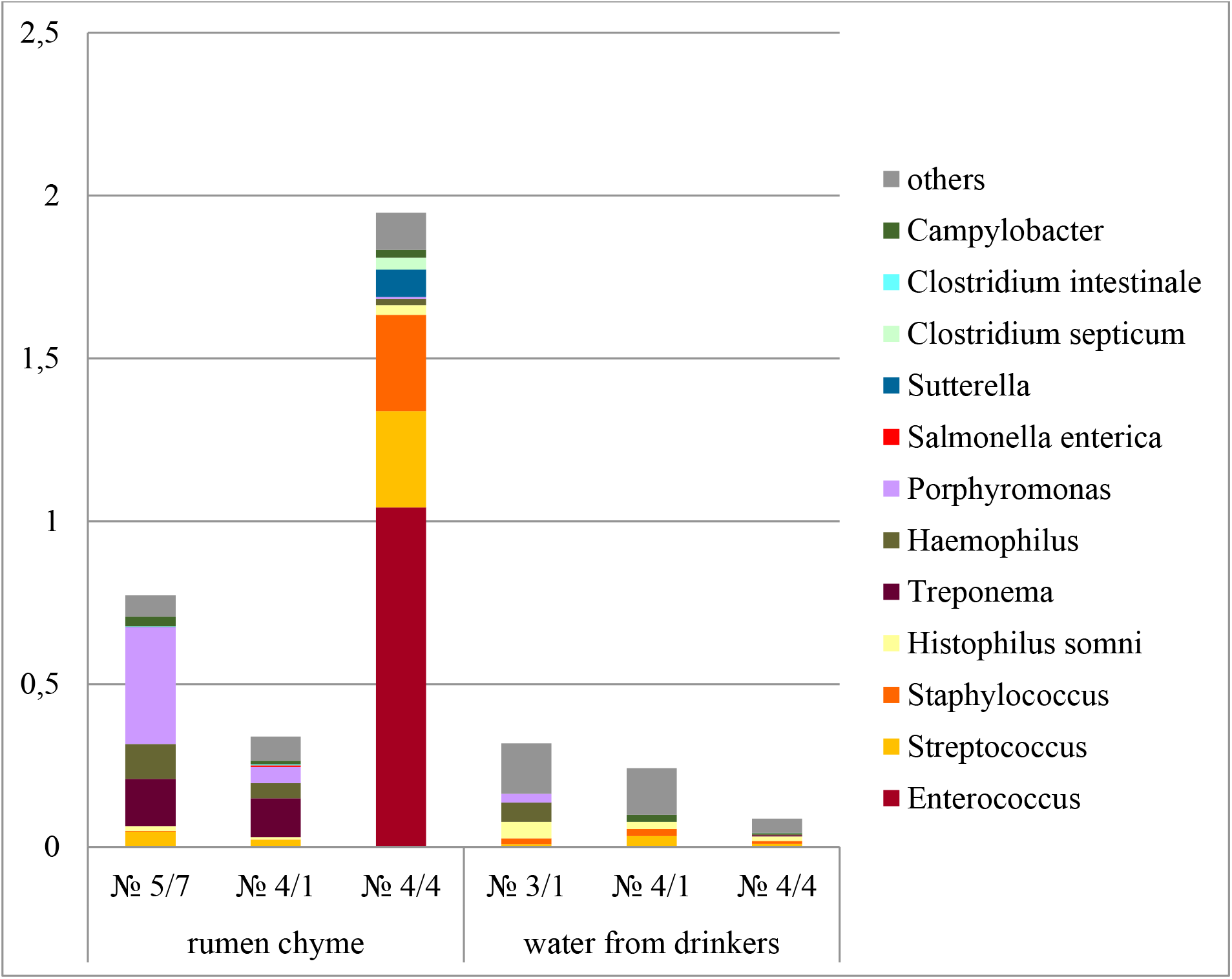
The content of all pathogenic bacteria in the samples, % (according to the analysis by the NGS-sequencing method), %.

Pathogenic species of *Porphyromonas* found are mainly associated with inflammation in the oral cavity. The largest proportion of pathogenic representatives of the genus *Porphyromonas* was found in a sample of cattle rumen fluid from building No. 3/1 - 0.4±0.03%. In other samples of cicatricial contents, the proportion of these bacteria was lower. Also, these bacteria were found in the water sample from the drinker of building No. 3/1 in the amount of 0.03±0.004%.

The proportion of pathogenic *Campylobacter* in samples of cicatricial contents ranged from 0.01±0.0006% (building No. 4/1) to 0.03±0.0009% (building No. 3/1). Also, these bacteria were found in water samples from the drinking bowls of buildings No. 4/1 in the amount of 0.02 ± 0.0015% and No. 4/4 in the amount of 0.002 ± 0.0001 %. Campylobacter bacteria can cause campylobacteriosis, an infectious disease of cattle livestock, manifested by lesions of the genital organs, frequent roaming, infertility, mass abortions and the birth of non-viable offspring.

Bacteria of the genus *Mycoplasma* can cause mycoplasmosis in livestock, a common infectious disease of cattle in the world, affecting adult cows and heifers, calves, and newborn young. With mycoplasmosis, animals develop mastitis, endometritis, vulvovaginitis, salpingitis, abortion, infertility, the birth of premature, underdeveloped calves incapable of life. The proportion of mycoplasmas in samples of cicatricial contents ranged from 0.018±0.0009% (building No. 4/4) to 0.044±0.0038% (building No. 4/1). Also, these bacteria were found in water samples from the drinkers of buildings No. 3/1 in the amount of 0.009 ± 0.0006% and No. 4/4 in the amount of 0.003 ± 0.0047%.

Also in some samples there were other representatives of pathogenic microflora, including: pathogenic clostridia (*Clostridium septicum, C. intestinale*) that can cause bacteremia, poisoning; *Salmonella enterica*, the causative agent of toxic infections; pathogenic treponemas, in cattle are mainly considered to be the causative agents of dermatitis, including ulcerative dermatitis of the udder; leptotrichia, causative agents of leptotrichosis of the genital tract, as well as fever, bacteremia; fusobacteria, causative agents of purulent-necrotic processes.

## Conclusions

Thus, the composition of microorganisms in samples of rumen fluid and water from drinking bowls was studied using the molecular genetic method of NGS-sequencing. The conducted studies showed differences in the microflora of the cicatricial contents of cows from different buildings. In the samples of the cicatricial contents of cows from buildings No. 3/1 and 4/1, the microbial landscape corresponded to the norm: a high content of representatives of normal microflora and a low content of pathogenic, opportunistic and undesirable types of microorganisms. At the same time, the samples of cicatricial contents from corps 4/4 differed from the samples from the other two corps. In samples from this farm building, up to 82±4.5% of the entire rumen microbiota was occupied by uncultivated bacteria. Among the identified microorganisms in the samples of cicatricial contents from cows from building No. 4/4, a low content of representatives of beneficial microflora was found: cellulolytic microorganisms, VFA-synthesizing microorganisms, while the highest among the studied samples were representatives of pathogenic, conditionally pathogenic, and also undesirable microflora. In particular, in this group, a high content of pathogenic enterococci, staphylococci, streptococci, Escherichia/Shigella was found.

In water samples, non-culturable bacteria also occupied the main share.

At the same time, in all samples, including water samples from drinking bowls, a variety of pathogenic microorganisms were found, including campylobacter, mycoplasma, pathogenic *Clotridia*, leptotrichia, *Salmonella* and others. Interestingly, a number of these microorganisms were also found in the rumen of cows. Previously [4] revealed *Salmonella* sp. in 0.8% of troughs and shigatoxigenic - *E. coli* O157 in 1.3% of troughs on the territory of livestock enterprises.

Our study has shown that drinkers can be a source of pathogenic bacteria exposure in cattle, including a number of foodborne pathogens, and this degree of bacterial contamination appears to affect the microbial composition of the rumen.

***The work was carried out within the framework of the state task with the financial support of the Ministry of Agriculture of the Russian Federation No. 1022041400153-7-2*.*8*.*1***

